# Structural insights into the viral proteins binding by TRIM7 reveal a general C-terminal glutamine recognition mechanism

**DOI:** 10.1101/2022.03.24.485560

**Authors:** Xiao Liang, Jun Xiao, Xuzichao Li, Yanan Wen, Xing Che, Yongjian Ma, Xingyan Zhang, Yi Zhang, Deng Jian, Peihui Wang, Chenghao Xuan, Guimei Yu, Long Li, Heng Zhang

**Affiliations:** Department of Biochemistry and Molecular Biology, School of Basic Medical Sciences, Tianjin Medical University, Tianjin 300070, China; Department of Immunology, School of Basic Medical Sciences, Tianjin Medical University, Tianjin 300070, China; Suzhou YDS Pharmatech Co., Ltd; Key Laboratory for Experimental Teratology of Ministry of Education and Advanced Medical Research Institute, Cheeloo College of Medicine, Shandong University, Jinan 250012, China

**Author notes:** Correspondence (G.Y.), (L.L.), (H.Z.). These two authors contribute equally to this work (Xiao Liang, Jun Xiao).

## Abstract

The E3 ligase TRIM7 has emerged as a critical player in viral infection and pathogenesis. A recent study found that TRIM7 inhibits human enteroviruses through ubiquitination and proteasomal degradation of viral 2BC protein by targeting the 2C moiety of 2BC protein. Here, we report the crystal structures of TRIM7 in complex with 2C, where the C-terminal region of 2C is inserted into a positively charged groove of the TRIM7 PRY-SPRY domain. Structure-guided biochemical studies revealed the C-terminus glutamine residue of 2C as the primary determinant for TRIM7 binding. Such a glutamine-end motif binding mechanism can be successfully extended to other substrates of TRIM7. More importantly, leveraged by this finding, we were able to identify norovirus and SARS-CoV-2 proteins, and physiological proteins, as new TRIM7 substrates. We further show that TRIM7 may function as a restriction factor to promote the degradation of the viral proteins of norovirus and SARS-CoV-2, thereby restoring the Type I interferon immune response and inhibiting viral infection. Several crystal structures of TRIM7 in complex with SARS-CoV-2 proteins are also determined, and a conserved C-terminus glutamine-specific interaction is observed. These findings unveil a common recognition mode by TRIM7, providing the foundation for further mechanistic characterization of antiviral and cellular functions of TRIM7.

## Introduction

The tripartite motif (TRIM) family proteins play essential roles in a plethora of cellular processes including signal transduction, oncogenesis, cell death and antimicrobial responses ^1,2^. TRIM proteins, belonging to the RING E3 ligase family, are characterized by the conserved RBCC domain architecture consisting of the RING (R), one or two B-boxes (B) and the coiled-coil (CC) domains, and a variable C-terminal domain (CTD)^3^. The RING domain recruits ubiquitin-loaded E2 and catalyzes the conjugation of ubiquitin to substrates, while B-box and coiled-coil domains mainly play regulatory roles. The CTD usually confers substrate specificity. More than ten distinct types of CTD have been identified, among which is the PRY-SPRY domain present only in vertebrates ^4,5^.

As a member of the TRIM family, TRIM7 is featured with one B-box domain in the RBCC architecture and a C-terminal PRY-SPRY domain. TRIM7 is implicated in multiple critical cellular processes, such as glycogen biosynthesis and oncogenic signaling pathways including c-Jun/AP1 activation and the DUSP6/p38 pathway through different substrates ^6–8^. Notably, TRIM7 is also emerging as a crucial player in viral pathogenesis. For example, a recent study has identified human TRIM7 as an intrinsic antiviral effector against human enteroviruses such as coxsackievirus B3 (CVB3) ^9^, which are positive-sense single-stranded RNA (+ssRNA) viruses. TRIM7-mediated ubiquitination and subsequent degradation of viral 2BC protein are implicated to inhibit viral infection. The non-structural protein 2C, which is derived from proteolysis of the precursor 2BC protein, functions as an ATPase/helicase to regulate viral RNA replication. TRIM7 recognizes the C-terminal region of 2C via PRY-SPRY domain, leading to ubiquitination and proteasomal degradation of viral 2BC protein but not 2C protein, thereby suppressing enterovirus replication ^9^. In addition, a genome-wide CRISPR screen revealed that TRIM7 also restricts norovirus infection in a ubiquitination-dependent manner through yet unknown substrates ^10^.

To elucidate the molecular mechanism of substrate recognition by TRIM7, we have determined multiple crystal structures of TRIM7 in complex with CVB3 2C protein. The structural studies combined with biochemical characterization allow us to propose a glutamine-end motif binding mechanism, where TRIM7 recognizes 2C mainly through C-terminus glutamine-specific interactions. Strikingly, this binding mode is found to be preserved for other targets. More importantly, leveraged by this new knowledge, we were able to predict and identify new viral targets in noroviruses and severe acute respiratory syndrome coronavirus 2 (SARS-CoV-2) virus, which are both +ssRNA viruses as with enteroviruses. Our findings not only unveil a general substrate binding principle for TRIM7, but also demonstrate a prevalent and versatile role of TRIM7 in +ssRNA virus infection and physiological processes such as tumorigenesis, potentially opening new avenues for therapeutic intervention in viral diseases and cancers.

## Results

### Biochemical characterization of TRIM7 and 2C association

Human TRIM7 restricts CVB3 enterovirus infection, by recognizing and degrading the viral protein 2BC, which could be further proteolytically processed into 2B and 2C proteins ^9^. The physical association was indicated to primarily occur between the PRY-SPRY domain of TRIM7 (hereinafter termed as TRIM7 unless otherwise indicated) and the 2C moiety of 2BC, especially the C-terminal region of 2C (117-329 aa, denoted as 2C unless otherwise stated) ^9^ (**Fig. 1a**). To further characterize the TRIM7-2C interaction, we purified the proteins and examined their binding using isothermal titration calorimetry (ITC). Indeed, TRIM7 tightly associated with 2C with a dissociation constant (*K*_D_) of ~ 6 *μ*M (**Fig. 1b**). By contrast, no obvious binding was observed after truncating the last 11 residues of 2C (117-318 aa), consistent with the previous pull-down assay results ^9^. Similarly, TRIM7 was stoichiometrically co-eluted with 2C but not the C-terminus deleted 2C protein in gel filtration (**Figs. 1c and S1a-c**). In line with these results, the C-terminus derived peptide fragment displayed a binding affinity of ~ 3 *μ*M toward TRIM7 (**Fig. S1d-e**), comparable to that of 2C protein, demonstrating a dominant role of this C-terminal motif in binding of TRIM7.

**Figure 1.**
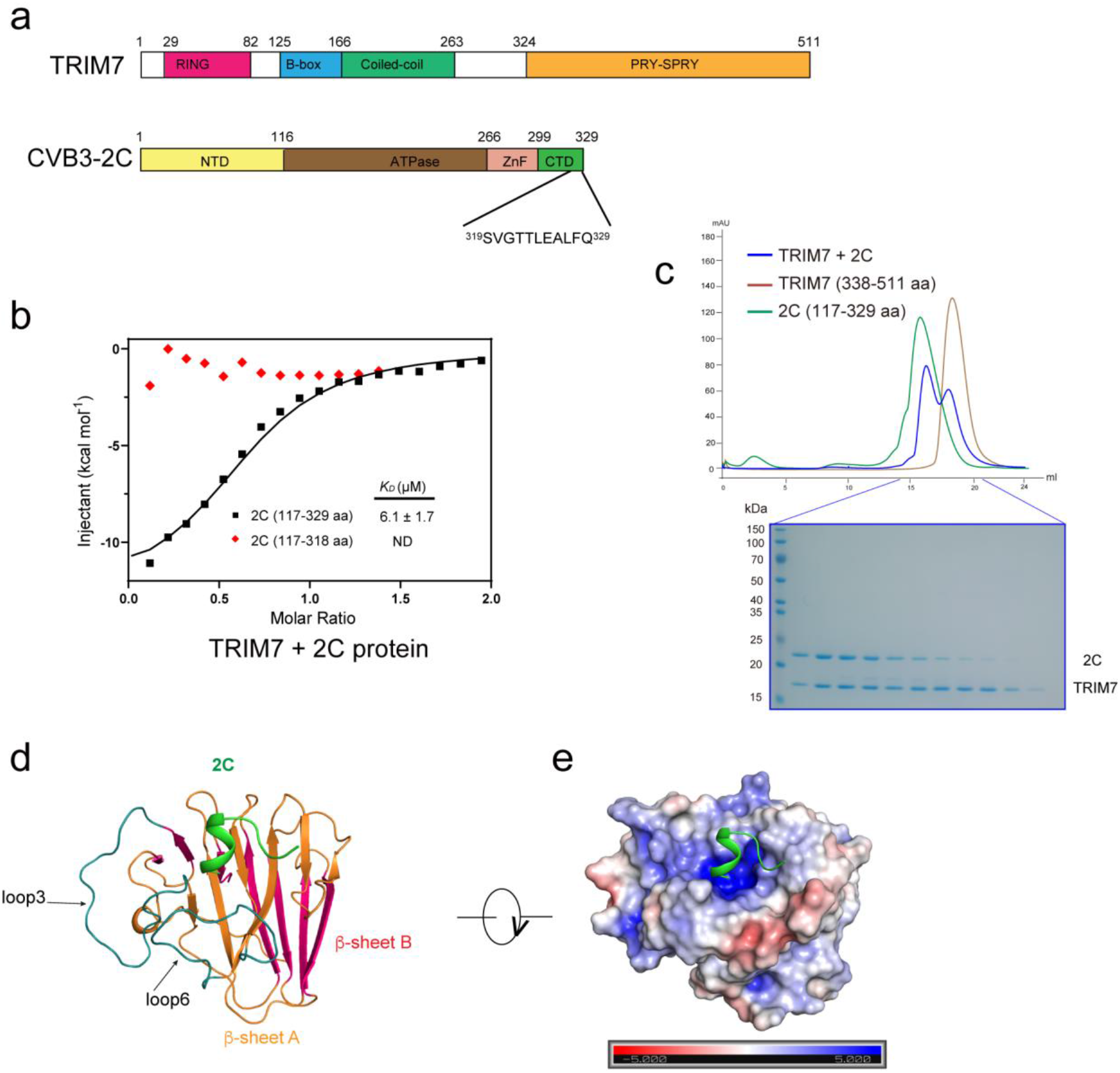
Biochemical and structural characterizations of TRIM7-2C complex. **a**, Domain organization of TRIM7 and 2C. NTD: N-terminal domain; ZnF: Zinc finger; CTD: C-terminal domain. **b**, Integrated heat plots for ITC measurements of 2C (in black) and C-terminus truncated 2C (in red) into TRIM7. ND, no detectable binding. **c**, Gel-filtration analysis of TRIM7 with 2C. Top panel: elution profile. Bottom panel: SDS-PAGE analysis of elution fractions. **d**, Crystal structure of TRIM7-2C complex. The bound 2C fragment is colored in green. TRIM7 PRY-SPRY domain is composed of two antiparallel *β* sheets (*β*-sheet A in orange, *β*-sheet B in violet red), connected with loops. Positions of loop3 and loop6 are indicated. **e**, The electrostatic potential of the TRIM7 PRY-SPRY domain.

Apart from the type B enterovirus CVB3, TRIM7 was also reported to inhibit other human enteroviruses such as enterovirus 71 (EV71, type A), poliovirus (PV, type C) and enterovirus D68 (EVD68, type D) ^9^. Sequence analysis of these 2C proteins revealed a highly conserved C-terminus (**Fig. S1d**), indicating that TRIM7 may use a general binding mode for different 2C proteins of human enteroviruses. This is supported by the results that TRIM7 binds to the C-termini of different 2C proteins with similar binding affinities (3-9 *μ*M) (**Fig. S1e**).

### Overall structure of the TRIM7-2C complex

Next, we set out to further elucidate the structural basis of TRIM7-2C complex association. Although extensive crystallization trials of TRIM7-2C complex failed, we were able to crystallize and determine multiple structures of TRIM7 (338-511 aa, PRY-SPRY domain) in complex with CVB3 2C-derived peptides (319-329 aa) at high resolutions (1.1-1.6 Å) (**Table S1**). Atomic protein models were built unambiguously for TRIM7 and the last four to ten residues of 2C (**Figs. 1d and S2a**). All the solved structures with different space groups are nearly identical (**Fig. S2b**), excluding the possibility of artifacts caused by crystal packing. TRIM7 exhibits a typical PRY-SPRY domain fold^4^, composed of two antiparallel *β*-sheets connected by loops of variable lengths (**Fig. 1d**). These two *β*-sheets, *β*-sheet A and *β*-sheet B, pack against each other in a twisted *β*-sandwich configuration with a concave surface at one side and a convex surface for the other side. The *β*-sheet A and loops 3 and 6 form a positively charged groove to accommodate 2C (**Fig. 1e**). Interestingly, this is different from a previously predicted binding mode between TRIM7 and glycogenin-1 (GN1)^11^. The binding affinity between TRIM7 and 2C was reduced by ~ 3-fold in a higher salt binding buffer, demonstrating that the polar contacts contribute to their physical association (**Fig. S2c**). Mapping of the evolutionary sequence conservation to TRIM7 structure revealed an invariant binding pocket (**Fig. S2d**), suggesting a conserved 2C recognition mode across TRIM7 orthologs.

### Interaction of TRIM7 with 2C

In the present structure, 2C displays a helical structure that inserts into the binding groove on the concave side of TRIM7. Specifically, the C-terminus Gln329 (termed as position −1) of 2C situated at the bottom of the binding pocket is involved in substantial interactions with TRIM7 (**Fig. 2a**). The side-chains of Asn383, Arg385 and Ser499 from TRIM7 make polar interactions with the carboxyl group of Gln329. On the other side, the amide moiety of Gln329 is buttressed by hydrogen bonds to the main-chains of Gly408 and Ser499, as well as the side-chains of Gln436 and Ser499 from TRIM7. The side-chain of Gln329 is further bracketed by Phe426 and Cys501 through hydrophobic interactions.

**Figure 2.**
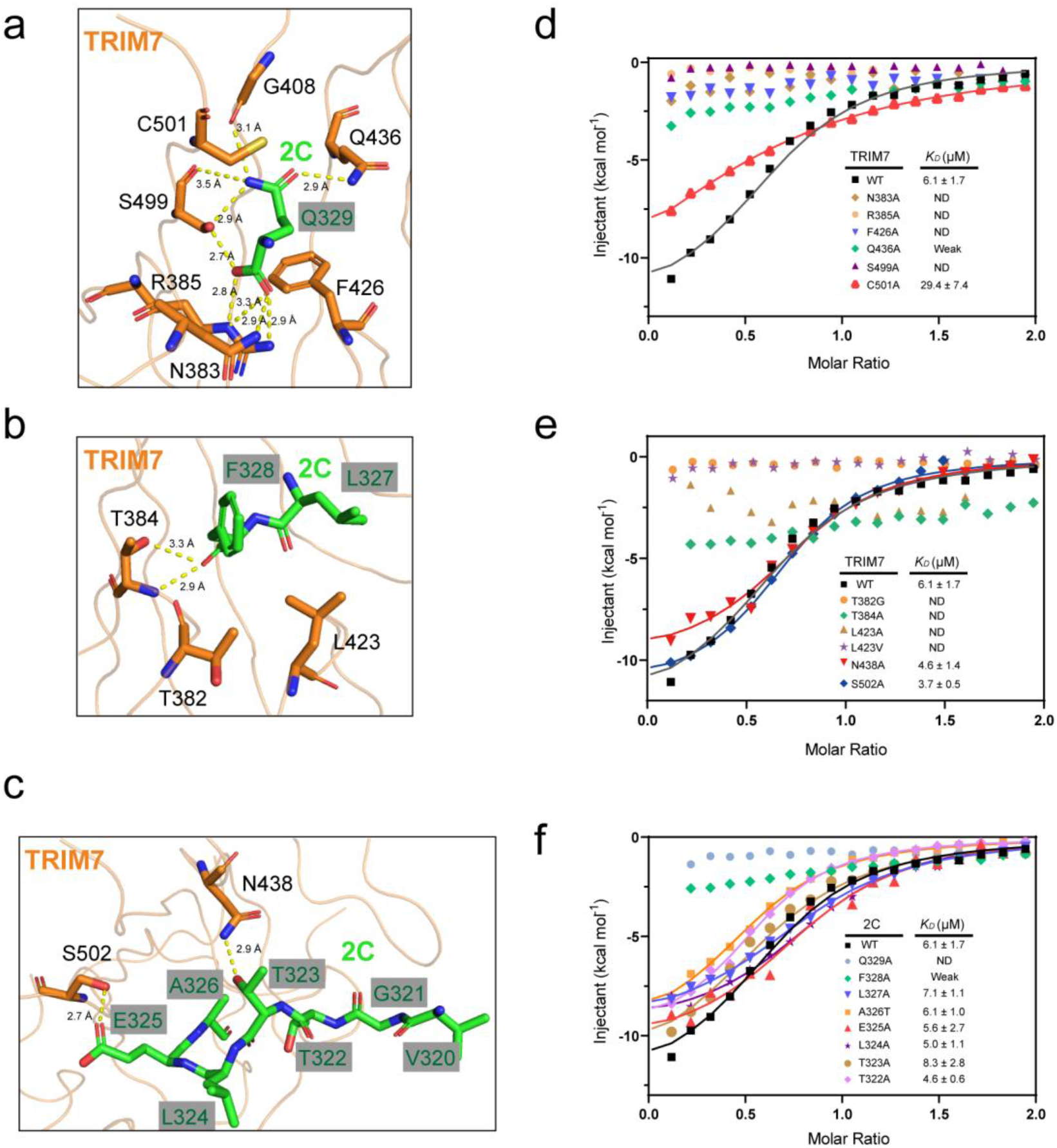
Interactions between TRIM7 and 2C. **a**, The C-terminus Q329 inserts into a pocket formed by residues N383, R385, G408, F426, Q436, S499 and C501 in TRIM7. **b**, The interaction interface between the two resides preceding Q329 in 2C (F328 and L327) and TRIM7. **c**, The interface between 2C residues 320-326 and TRIM7. **d-e**, ITC measurements of TRIM7 mutants with 2C. **f**, Alanine-scanning mutagenesis of 2C. Integrated heat plots for ITC measurements of 2C mutants to TRIM7 were displayed.

The carbonyl group of Phe328 (position −2) of 2C engages in hydrogen bonds with Thr384 of TRIM7 (**Fig. 2b**), while its aromatic ring is further stabilized by hydrophobic contacts with Thr382 and Leu423. Leu327 (position −3) of 2C is anchored in place mainly through the hydrophobic interaction with Leu423 of TRIM7. Nevertheless, few contacts were observed beyond the last three residues of 2C (**Fig. 2c**).

### Mutagenesis studies of the TRIM7-2C binding interface

To explore the importance of the observed interactions, we introduced point mutations into TRIM7 and measured their binding affinities. Replacement of the Gln329-interacting residues with alanine, including Asn383, Arg385, Phe426, Gln436, Ser499 and Cys501, significantly reduced or abolished 2C binding (**Fig. 2d**). Similarly, the interaction of TRIM7 with 2C was disrupted when mutating the Phe328-interacting residues (Thr382, Thr384 and Leu423) (**Fig. 2e**). However, substitution of other residues (such as Asn438 and Ser502) in the binding groove with alanine had no or very little effect on the binding affinity (**Fig. 2e**).

To further discern the residues of 2C crucial for TRIM7 binding, we generated point mutations on 2C for binding studies. Not surprisingly, both the Q329A and F328A mutations significantly impaired the binding (**Fig. 2f**). Nonetheless, alanine substitutions of Leu327-Thr322 had no impact on binding. These results demonstrated the critical role of the last two residues of 2C, Gln329 and Phe328, in binding of TRIM7.

### TRIM7 specifically recognizes the C-terminus glutamine-containing motif

As aforementioned, the last two or three residues in 2C seem to be the determinant for TRIM7 binding. To further understand the binding preference of TRIM7, we generated a series of 2C mutants on the last three positions and examined their binding. Gln329 at position −1 fits snugly into the binding pocket (**Fig. 3a**). Substitution of Gln329 with bulkier residues, such as tyrosine, arginine and methionine, that is expected to have steric clashes, exhibited no binding to TRIM7 (**Fig. 3b**). Likewise, replacement of Gln329 with smaller residues (alanine, threonine, asparagine and valine) disrupted the TRIM7 binding, likely due to loss of the amide group-mediated interactions with Gln436 and Ser499. Notably, mutation of Gln329 to glutamate with a highly similar chemical structure also abrogated the binding. Molecular dynamics (MD) analysis revealed that Q329E mutation affects the solvation and interrupts hydrogen-bonding with TRIM7 (**Figs. 3c and S3a and Table S2**), which could explain the profoundly weakened binding. The free C-terminal carboxyl group is also critical for TRIM7 binding, which is supported by the finding that no detectable binding was observed in the presence of the C-terminal amidation modification (**Fig. 3b**). Collectively, these results suggest that a glutamine residue at position −1 is indispensable and irreplaceable for TRIM7 binding.

**Figure 3.**
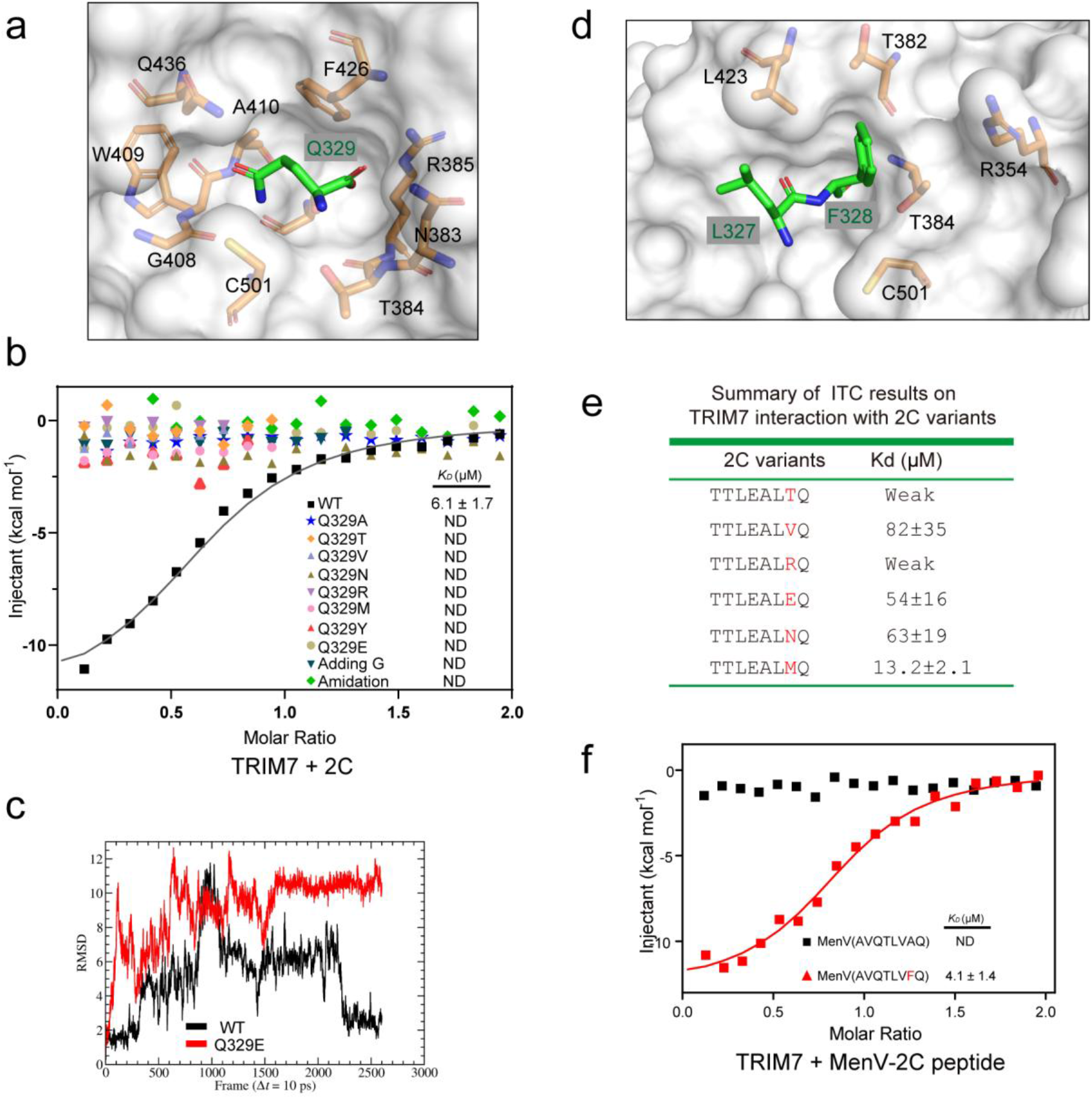
Recognition mechanism of 2C by TRIM7. **a**, The C-terminus glutamine-binding pocket in TRIM7. Q329 (C-terminus, −1 position) of 2C is colored in green. **b**, Replacement or manipulation of Q329 at −1 position blocked TRIM7 binding, as revealed by the ITC measurements. **c**, MD simulation of Q329E mutation for TRIM7-2C complex formation. **d**, The pocket in TRIM7 accommodating F328 (−2 position) and L327 (−3 position) of 2C. **e**, ITC measurements of 2C mutants at the −2 position, to TRIM7. **f**, Integrated heat plots for ITC measurements of TRIM7 with peptide fragments of 2C protein in MenV non-human enterovirus.

Phe328 at position −2 rests against the rim of the pocket with its aromatic ring surrounded by the hydrophobic side-chains of Leu423, Thr382 and Thr384 (**Fig. 3d**). Replacement of Phe328 with smaller residues (alanine, threonine, valine, asparagine, glutamate), with the potential to lose the phenyl group-mediated hydrophobic interaction, substantially reduced TRIM7 binding (**Figs. 2f and 3e**). Arginine substitution of Phe328 would also weaken the interaction due to the electrostatic repulsion with Arg354 of TRIM7, which agrees with our finding that F328R mutation significantly diminished the binding of TRIM7. However, substitution of Phe328 with bulkier residues such as methionine and tyrosine would be better tolerated. Indeed, F328M had a similar binding affinity to TRIM7 as WT (**Fig. 3e**). Leu327 at position −3 protrudes outwards from the pocket (**Fig. 3d**), forming loose interactions with the binding surface of TRIM7. Different types of amino acid substitutions at position −3 were all tolerated (**Fig. S3b**). Taken together, our results demonstrate that TRIM7 not only specifically recognizes glutamine at position −1, but also has a preference for bulky hydrophobic residues at position −2. This observation could explain why 2C protein from non-human enterovirus mengovirus (MenV), which harbors an alanine residue at the position equivalent to Phe328 in CVB3 2C protein, fails to interact with TRIM7 ^9^. Mutating alanine to phenylalanine in 2C of MenV was sufficient to restore the TRIM7 binding (**Fig. 3f**), highlighting the importance of the secondary determinant involving −2 position.

### Identification of the potential norovirus substrate for TRIM7

TRIM7 was recently reported to inhibit the replication of norovirus ^10^. However, the target of TRIM7 remains poorly understood. Referring to the observed glutamine-end motif binding rule, we therefore examined the sequence information of norovirus proteins. Several viral proteins, such as capsid protein VP1, p48, NTPase and the 3C-like protease, were found bearing the TRIM7 recognition motif, among which NTPase was found most conserved in different genogroups (GI-GV) (**Fig. 4a**). We therefore focused our study on NTPase.

**Figure 4.**
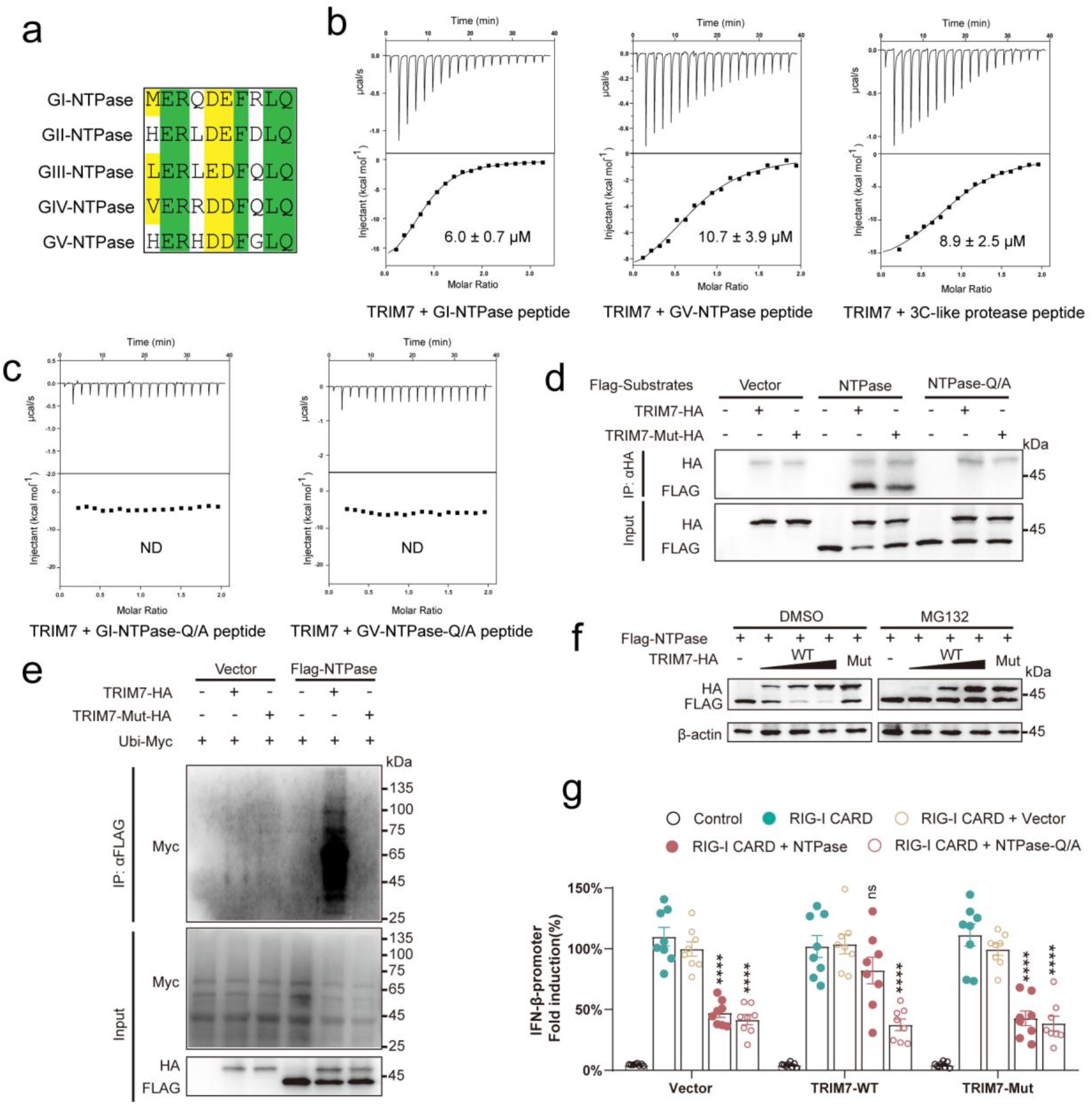
TRIM7 binds and ubiquitinates norovirus NTPase protein. **a**, multiple sequence alignment of norovirus NTPase proteins from different genogroups. The aligned C-terminal last 10 residues were shown. The GI-GIV NTPase proteins are derived from human noroviruses and the GV NTPase protein originates from murine norovirus. **b**, Thermodynamic analysis of the interaction between TRIM7 and NTPase peptide fragments from GI (left panel) and GV (middle panel), and 3C-like protease (right panel). **c**, The C-terminus glutamine mutation in GI or GV NTPase abolished the TRIM7 binding as determined by ITC. Q/A, alanine replacement of the C-terminus glutamine. **d**, Human norovirus NTPase co-immunoprecipitated with TRIM7. Flag-NTPase or empty vector was co-transfected with TRIM7-HA in HEK293T cells. TRIM7-Mut-HA, TRIM7 catalytic dead mutant (C29A/C31A); NTPase-Q/A, the alanine replacement of the C-terminal glutamine in NTPase. **e**, Expression of TRIM7 caused ubiquitination of human norovirus NTPase. Flag-NTPase or empty vector was co-transfected with Ubi-Myc in the presence or absence of TRIM7-HA in HEK293T cells. **f**, TRIM7 promoted the NTPase degradation in a dose-dependent fashion. Cellular degradation assays in cells expressing the TRIM7 and NTPase proteins indicated. Cells were treated with DMSO or MG132. Mut, TRIM7 catalytic dead mutant. **g**, IFN-β reporter assay. HEK293T cells were transfected with an IFN-β promoter-driven luciferase reporter construct, a RIG-I CARD expression plasmid and the indicated NTPase expression plasmid in the presence or absence of TRIM7. Relative luciferase activity was quantified 24 h post-transfection. Error bars represent mean ± SEM of technical triplicates. Statistical significance was determined by comparing each respective “RIG-I 2C + Vector” group using two-way ANOVA with Dunnett’s correction, ****p < 0.0001. ns, not significant.

ITC measurement showed that the NTPase C-terminal peptide fragments from both human (GI) and murine (GV) noroviruses directly interact with TRIM7 with *K*_D_ values of ~ 6-10 μM (**Fig. 4b**), while alanine replacement of the C-terminus glutamine in either GI or GV NTPase abolished the TRIM7 binding (**Fig. 4c**). Consistently, WT NTPase protein derived from human norovirus, but not the C-terminal alanine-substituted NTPase protein, co-immunoprecipitated with TRIM7 (**Fig. 4d**). These results demonstrate that TRIM7 indeed can target norovirus NTPase protein using the glutamine-end motif binding mechanism both in vitro and in vivo.

We next asked whether TRIM7 could ubiquitinate and degrade NTPase. Either WT TRIM7 or ligase dead TRIM7 (C29A/C32A) was co-transfected with NTPase from human norovirus and ubiquitin into the HEK293T cells. As expected, WT TRIM7, but not the ligase dead mutant, robustly ubiquitinated NTPase (**Fig. 4e**). Furthermore, expression of WT TRIM7 triggered the degradation of NTPase, whereas the degradation was inhibited by treatment with the proteasome inhibitor MG132 but not the DMSO (**Fig. 4f**), implicating the 26S proteasome-dependent degradation. The NTPase protein possesses fundamental enzymatic activities required for viral replication ^12^. It has been demonstrated that norovirus NTPase impairs the Type I interferon-β (IFN-β) expression ^13^. We next determined whether TRIM7 interferes with this NTPase-mediated suppression. To this end, NTPase expression plasmid was co-transfected with the IFN-β luciferase reporter plasmid and the vectors expressing RIG-I CARD and TRIM7. The expression of NTPase inhibited the IFN-β production, whereas WT TRIM7 restored the expression level of IFN-β (**Fig. 4g**), suggesting a restriction role of TRIM7 in norovirus infection. However, the ligase dead TRIM7 mutant had no obvious effect on IFN-β expression. Similarly, the NTPase-mediated suppression of IFN-β was not affected regardless of the presence or absence of WT TRIM7 when the C-terminus glutamine of NTPase was mutated to alanine (**Fig. 4g**), further demonstrating the importance of C-terminal glutamine-specific recognition. Hence, it is likely that TRIM7 may inhibit norovirus infection through degradation of the NTPase protein.

### TRIM7 recognizes and ubiquitinates SARS-CoV-2 proteins

As TRIM7 restricts the infection of human enterovirus and norovirus, both of which are +ssRNA viruses, it raises an interesting question whether TRIM7 could function as a broad-spectrum regulator in +ssRNA virus infection. We therefore attempted to explore whether another +ssRNA virus, SARS-CoV-2, could be targeted by TRIM7 based on the identified substrate recognition rule. Sequence analysis was performed for proteins encoded by SARS-CoV-2 genome, surprisingly revealing that most of the nonstructural proteins (NSP) including NSP4-10, NSP12-15, and the structural membrane (M) protein have the glutamine-end motif (**Fig. 5a**). ITC experiments were then performed to evaluate their association with TRIM7. As revealed, binding was detected for all the measured peptides derived from the C-termini of these candidates (**Fig. 5b**). Supporting our observation, a recent proteomics study also shows that TRIM7 interacts with the M protein of SARS-CoV-2 ^14^. To further confirm the interaction, full-length NSP5 protein was purified, which consistently was found bound with TRIM7 (**Fig. 5c**). These were in agreement with the co-IP experiment, where WT NSP5 and NSP8, but not the alanine mutations at the C-terminal glutamine, were detected in the immunoprecipitated samples of TRIM7 (**Fig. S4a**). Furthermore, expression of TRIM7 triggered the ubiquitination and proteasomal degradation of NSP5 and NSP8 (**Figs. 5d-f**). Together, these results demonstrate that TRIM7 could target a large array of SARS-CoV-2 viral proteins, which may modulate the SARS-CoV-2 infection and pathogenesis. We further determined the crystal structures of TRIM7 in complex with NSP5-, NSP8- and NSP12-derived peptides at high resolutions (1.2-1.4 Å) (**Figs. 4g-h and S4b**) (**Table S3**). Although the C-terminal sequences among these three NSP proteins are divergent (**Fig. 5a**), their binding modes are similar to that of 2C in TRIM7-2C complex, suggesting a conserved C-terminal glutamine recognition mechanism.

**Figure 5.**
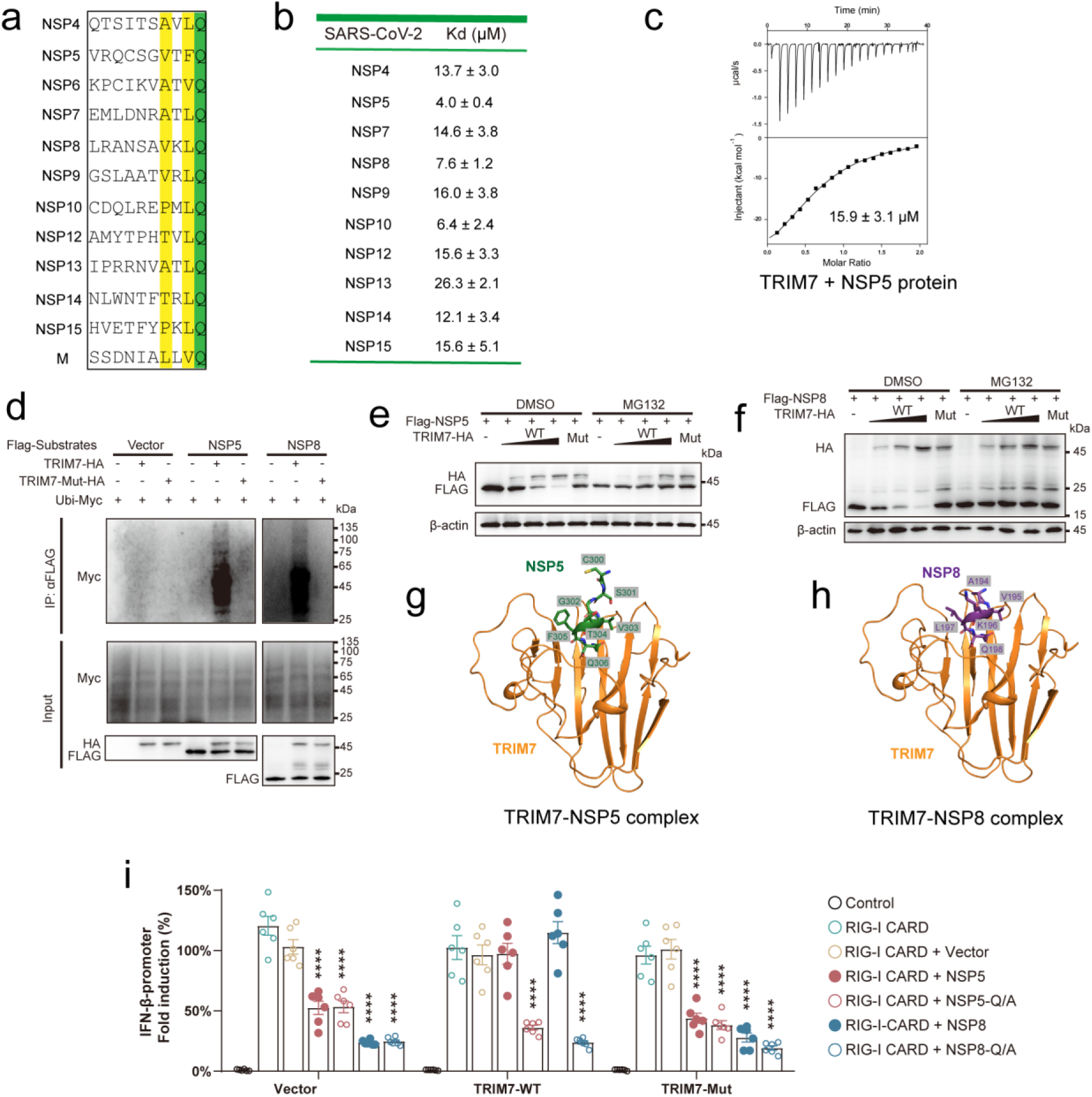
TRIM7 targets a large fraction of viral proteins in SARS-CoV-2. **a**, Multiple sequence alignment of SARS-CoV-2 viral proteins ended with glutamine. The aligned C-terminal last 10 residues were shown. **b**, Thermodynamic analysis of the interaction between TRIM7 and peptides derived from SARS-CoV-2 proteins. **c**, ITC measurement of purified full-length NSP5 of SARS-CoV-2 to TRIM7. **d**, In vivo ubiquitination assay for NSP5 and NSP8. Flag-tagged NSP5, NSP8 or empty vector were co-transfected with Ubi-Myc in the presence or absence of TRIM7-HA in HEK293T cells. TRIM7-Mut-HA, TRIM7 catalytic dead mutant (C29A/C31A). **e-f**, TRIM7 induced the proteasomal degradation of NSP5 (e) and NSP8 (f) in a dose-dependent fashion. Cellular degradation assays in cells expressing the TRIM7 and NSP proteins indicated. Cells were treated with DMSO or MG132. Mut, TRIM7 catalytic dead mutant. **g-h**, Overall structures of TRIM7 in complex with NSP5 (residues 300-306) (g) or NSP8 (residues 194-198) (h). **i**, IFN-β stimulation determined by a luciferase reporter. HEK293T cells were transfected with a luciferase reporter plasmid under the control of the IFN-β promoter, a RIG-I CARD plasmid and the indicated NSP proteins expression plasmid along with or without the TRIM7 expression plasmid. Relative luciferase activity was quantified 24 h post-transfection. Error bars represent mean ± SEM of technical triplicates. Statistical significance was determined by comparing each respective “RIG-I 2C + Vector” group using two-way ANOVA with Dunnett’s correction, ****p < 0.0001.

We next sought to characterize the role of TRIM7 in SARS-CoV-2 viral infection. Given that SARS-CoV-2 proteins could antagonize IFN signaling ^15–20^, we co-transfected HEK293T cells with the plasmid encoding individual SARS-CoV-2 NSP proteins, a IFN-β luciferase reporter plasmid and a RIG-I CARD expression plasmid. Consistent with recent studies, we observed that expression of SARS-CoV-2 viral proteins including NSP5, NSP6 and NSP8 significantly suppressed IFN-β production (**Fig. S4c**), suggesting that these NSP proteins may inhibit IFN signaling to evade host immune response. However, IFN-β expression was rescued in the presence of WT TRIM7 but not the ligase dead mutant, indicative of TRIM7 as a restriction factor in viral infection (**Fig. 5i**). We next explored how these NSP proteins affect IFN response. TBK1 phosphorylation, a key event for IFN signaling, was impaired when expressing NSP5 or NSP8 protein (**Fig. S4d-e**). As expected, WT TRIM7, but not the catalytic dead TRIM7 mutant, compromised the inhibition effects of NSP5 and NSP8 proteins. To reconcile the glutamine-end binding mode, we also introduced mutation into the C-terminal glutamine residues in NSP5 and NSP8. These mutations blocked the IFN-β production and TBK1 phosphorylation similar to the WT NSP5 and NSP8 (**Figs. 5i and S4d-e**). Nonetheless, TRIM7 failed to restore IFN response in the presence of these NSP mutations. Together, our data demonstrate that TRIM7 may function as a restriction factor in SARS-CoV-2 infection through glutamine-end specific recognition mode.

### Recognition of cellular proteins by TRIM7

TRIM7 also regulates diverse human proteins to exert its cellular functions. We next asked whether the same recognition mode of viral proteins by TRIM7 is also adopted by the cellular targets. Multiple cellular proteins that have been previously reported as putative targets of TRIM7 were explored, including BRMS1, DUSP6, GN1, RACO-1 and STING ^7,8,21–23^. Indeed, TRIM7 was able to directly interact with the C-terminal peptide fragments of GN1 (~ 8 μM) and RACO-1 (~ 10 μM) containing tolerable glutamine-end motifs (**Fig. S5a**). Mutation of the C-terminus glutamine in RACO-1 completely abolished TRIM7 binding (data not shown).

In contrast, the peptide or protein fragments, which are derived from the C-termini of BRMS1, DUSP6 and STING without glutamine-end, exhibited no obvious binding toward TRIM7 (**Fig. S5b**). The potential explanation is that these proteins could possibly interact with TRIM7 either through additional binding partners or a different interface ^24,25^. Taken together, these results indicate that the glutamine-end motif binding mechanism is a common, albeit potentially not the only, target recognition strategy of TRIM7 for both human and viral proteins.

TRIM7 has emerged as an essential regulator of cancer development ^7,8,21,26,27^. Several cancer-related TRIM7 somatic mutations in the COSMIC database (N383S, R385C, L423P, S502F) ^28^, which are located in the substrate binding pocket (**Figs. 2a-c and S5c**), failed to bind the substrate in our ITC experiments (**Fig. S5d**). It will be interesting to further investigate these mutations in the context of oncogenesis.

## Discussion

TRIM7, a RING type E3 ligase, is implicated in viral infection as well as oncogenesis ^7–10,21^. However, the detailed mechanism by which TRIM7 recognizes substrate is unclear. In this study, we structurally and biochemically characterized the interaction between TRIM7 and different viral substrates, including protein 2C of CVB3 enterovirus, and non-structural proteins of SARS-CoV-2, revealing a common target binding mode, thereby enabling the identification and discovery of new substrates, such as viral proteins in positive sense single-stranded RNA (+ssRNA) viruses.

### The target recognition mechanism of TRIM7

The C-terminus Gln329 of 2C protein (position −1) is buried in the binding pocket of TRIM7, establishing extensive polar and hydrophobic contacts (**Fig. 2a**). Our data demonstrate the glutamine-end as the primary determinant for TRIM7 binding. TRIM7 also displays a strong preference for bulky hydrophobic amino acids at position −2. This explains why the MenV 2C protein terminating with glutamine (alanine at position −2) fails to interact with TRIM7 ^9^ (**Fig. 3f**). Therefore, TRIM7 has best preference for substrate with the C-terminal φQ (φ denotes bulky hydrophobic residue) motif. However, the other positions of substrate may also affect the TRIM7 binding, especially in the cellular context. It is possible that TRIM7 might be able to bind the proteins harboring the unfavorable C-terminal motif such as AQ under the physiological conditions, albeit with low binding affinity in our assay. Future studies would be required to systematically investigate the TRIM7 binding in the cellular context.

### Prevalent and versatile roles of TRIM7 in +ssRNA virus infection

TRIM7 was reported to suppress human enterovirus replication by degrading viral protein 2BC but not 2C, although the 2C part in 2BC is responsible for TRIM7 binding ^9^. It is likely that a precise orientation of substrate would be required for the ubiquitin deposition ^29^. As previously reported, alanine mutation of Thr323 at 2C would confer resistance to TRIM7 ^9^. In our structure, the hydroxyl group of Thr323 is hydrogen bonded to Asn438 of TRIM7 (**Fig. 2c**). However, mutations of Thr323 including T323A, T323G, T323S and T323I had little effect on the binding (**Fig. S6**). One proposed possibility is that these variants would alter the plasticity and enzymatic activity of 2C to gain a fitness advantage in vivo ^9^. The N-terminal domain induces the hexamerization of 2C. Alternatively, these mutations may affect the binding in the context of full-length rather than the N-terminus deleted 2C protein.

We also identified and demonstrated the NTPase protein, which harbors a highly conserved glutamine-end across norovirus genogroups, as a substrate of TRIM7. Ubiquitination and proteasomal degradation of norovirus NTPase could potentially suppress the enzymatic activities necessary for viral replication ^12^, and counteract its negative effect on IFN-β production ^13^ (**Fig. 4g**), which thereby inhibits norovirus infection ^10^. In addition, the 3C-like protease possessing a C-terminus glutamine residue in murine norovirus but not in other noroviruses could be an extra, murine norovirus-specific TRIM7 target (**Fig. 4b**).

Our study further indicates TRIM7 as a potentially essential restriction factor for SARS-CoV-2 infection. Through the same glutamine-end motif recognition mechanism, TRIM7 targets the structural protein M, nonstructural proteins NSP4-10 and NSP12-15, which are all essential proteins for SARS-CoV-2 viral replication ^30^. Ubiquitination and subsequent degradation of these NSP proteins by TRIM7 could affect their functions such as suppression of immune signaling (**Fig. 5d-f,i**). Intriguingly, TRIM7 could readily target all the components of the replication-transcription complex composed of NSP7, 8, 12 and 13 ^31^ (**Fig. 5b**), which are promising targets for COVID19 disease treatment ^32,33^. It is therefore likely that the TRIM7-triggered ubiquitination and proteasomal degradation, or sequestration of these components through binding could possibly lead to suppression of the SARS-CoV-2 replication. However, further studies will be needed to explore the relationship between TRIM7 and SARS-CoV-2 infection.

TRIM7 has also been reported to play a positive role in Toll-like receptor-mediated innate immune response that could be triggered by ssRNA virus infection ^34^. Collectively, these findings imply a potentially prevalent role of TRIM7 E3 ligase in restricting +ssRNA virus infection. Interestingly, TRIM7 was also found involved in promoting Zika virus (+ssRNA) entry by ubiquitinating the viral envelope protein ^35^, indicating potential functional versatility of TRIM7 in response to different +ssRNA virus infections.

### Cleavage specificity of viral protease determines the vulnerability to TRIM7?

The +ssRNA viral genomes generally encode polyproteins, which are proteolytically cleaved into individual structural and nonstructural viral proteins by viral proteases. The substrate cleavage specificity of these viral proteases therefore determines the C-terminal sequences of the mature viral proteins. The 3C or 3C-like proteases of human enterovirus, norovirus and SARS-CoV-2 virus show cleavage preference for the glutamine at the position P1 ^36–39^, releasing glutamine-terminated viral proteins, which explains the vulnerability of their mature viral proteins to TRIM7 targeting and inhibition. Examination of more viral proteases thus may further extend the spectrum of +ssRNA viruses targeted by TRIM7.

### Identification and prediction of potential substrate

Several proteins have been reported as putative cellular substrates of TRIM7 ^6–8,21,27^, among which direct binding was detected for those possessing the glutamine-end motif but not others. One possible explanation is that TRIM7 may interact with bridging proteins to exert E3 ligase activity. Alternatively, other regions may be involved in substrate binding. Such a mechanism can be supported by the recent observations that the coiled-coil domain of TRIM7 is responsible for binding of MAVS protein ^25^. As there are several thousands of human proteins terminated with glutamine, the recognition principle described here would enable the identification of more physiological targets of TRIM7.

### A Gln/C-degron pathway?

The ubiquitin-proteasome system (UPS) controls protein stability through recognizing degradation signals, which is generally a motif of fewer than ten residues in the N- or C-terminus, namely N- or C-degrons, respectively ^40^. Several classes of C-terminal degrons such as Arg/C-end, Glu/C-end, Gly/C-end and Ala/C-end have been discovered ^41,42^. The observed C-end glutamine rule by the TRIM7 E3 ligase in UPS may represent a new type of Gln/C-end degron pathway.

In summary, our study has elucidated a general binding principle for both viral and physiological substrates of TRIM7, which would guide further biological studies of TRIM7 functions and the development of antiviral and disease therapeutics.

## Materials and Methods

### Protein expression and purification

DNA encoding TRIM7 (338-511 aa) and CVB3 2C (117-329 aa) protein were synthesized (Genewiz), and cloned into expression vectors (pET15 for TRIM7, pET28-SUMO for 2C). Point mutations and truncations were generated using the PCR site-directed Quick-Change mutagenesis. The resultant plasmids were transformed into competent BL21(DE3) *E. coli* cells for protein expression. Cells were grown to mid-log phase and 0.2 mM isopropyl-β-d-thiogalactoside (IPTG) was added to induce protein expression. Pelleted cells were resuspended in the lysis buffer A (25 mM Tris-HCl pH 8.0, 500 mM NaCl, 10 mM imidazole, 2 mM β-mercaptoethanol) and lysed by sonication. Ni-NTA resin (QIAGEN) was added to the lysate, incubated at 4°C for 1 h and then washed with buffer A. Protein was eluted with the elution buffer (25 mM Tris-HCl pH 8.0, 500 mM NaCl, 300 mM imidazole, 2 mM β-mercaptoethanol). His-tagged TRIM7 was further purified using the Superdex 200 size exclusion column with a running buffer containing 25 mM Tris-HCl pH 8.0, 150 mM NaCl, 2 mM DTT. TEV was used to remove the His-SUMO tag for 2C, which was further purified by ion-exchange chromatography using HiTrap Q column (Cytiva). Peak fractions were collected and concentrated for use.

We also generated a fusion construct in which 2C (319-329 aa) was C-terminally linked to TRIM7 (338-511aa) using a (GSA)_5_ linker. The fusion protein was purified by the same protocol as above. DNA encoding peptides derived from the C-terminal last 10 residues of the norovirus and cellular proteins were cloned into the pET-His-GST vector (Addgene:29655) with an N-terminal GST tag. Proteins were purified using the GST affinity chromatography (Cytiva). The eluted proteins were further purified using the HiTrap Q column. Peak fractions were collected and concentrated for the ITC binding experiments. The plasmid encoding NSP5 was a gift from Dr. Jian Lei. Protein expression and purification was performed as previously described ^43^.

### Crystallization, data collection and structure determination

TRIM7 was concentrated to ~16 mg/ml and incubated with synthesized peptides (Sangon Biotech) on ice for 30 min with a molar ratio of 1:2. The fusion protein was concentrated to ~15 mg/ml. Crystallization was performed using the sitting-drop vapor diffusion method at 18°C. The crystals were obtained by mixing 1 μl protein solution and 1 μl crystallization buffer. The TRIM7-2C peptide complex crystals were grown in a reservoir solution containing 100 mM citric acid pH 5.0, 20% (w/v) PEG6000, or 20 mM potassium nitrate, 20% (w/v) PEG3350. The TRIM7-2C fusion protein crystal was grown in a reservoir solution containing 200 mM ammonium acetate, 150 mM magnesium acetate tetrahydrate, 5% (w/v) PEG4000. The TRIM7-NSP5 complex crystals were grown in a reservoir solution containing 1.2 M potassium sodium tartrate tetrahydrate, 100 mM Tris pH 8.0. The TRIM7-NSP8 complex crystals were grown in a reservoir solution containing 2.5 M sodium chloride, 100 mM sodium acetate/ acetic acid pH 4.5, 200 mM lithium sulfate. The TRIM7-NSP12 complex crystals were grown in a reservoir solution containing 200 mM DL-malic acid pH, 20% (w/v) PEG3350.

Crystals were cryoprotected in the mother liquor supplemented with 20% glycerol before flash frozen in liquid nitrogen. X-ray diffraction datasets were collected at beamline BL18U1 of Shanghai Synchrotron Radiation Facility (SSRF) ^44^. Structures were solved by molecular replacement using TRIM7 structure (PDB ID: 6UMA) as a searching model in CCP4I2 package^45^. Structures of bound peptides were manually constructed using the COOT program ^46^. The refinement of structures was performed with PHENIX ^47,48^ and CCP4I2 ^45^. Statistics of data processing and structural refinement are listed in Table S1 and S3.

### Isothermal titration calorimetry

All the isothermal titration calorimetry (ITC) experiments were carried out in a buffer containing 25 mM Tris-HCl pH 8.0, 150 mM NaCl (unless otherwise stated) using a MicroCal PEAQ-ITC system (Malvern) at 25°C. A typical titration experiment involved 19 injections of protein or peptide substrate (300 μM to 2 mM) solution into the cell containing TRIM7 protein (20-50 μM). Data was analyzed using MicroCal PEAQ-ITC analysis software.

### Gel-filtration assay

Superdex 200 Increase 10/300 column (Cytiva) was used to carry out the gel-filtration assay. TRIM7 and 2C proteins were incubated on ice for 30 min at a 3:1 molar ratio before subjected to the column pre-equilibrated with the running buffer (25 mM Tris-HCl pH 8.0, 150 mM NaCl, 2 mM DTT). Peak fractions were collected and analyzed using SDS-PAGE.

### Molecular dynamics simulation

To understand the effect of the Q329E mutation to the TRIM7-2C complex, molecular dynamics (MD) simulation was performed. Amber FF14SB force field ^49^ was used for the wild type and Q329E complexes. Both complexes are solvated in TIP3P water so that the minimum distance between the box boundary and any atom in the complexes is 10 Å. As a result, 11182 and 11179 water molecules are added to the complex systems, correspondingly. Both systems follow the same simulation routine. The time step is 2 fs with the SHAKE algorithm ^50^ to fix the hydrogen atoms. The cutoff for calculating the electrostatic interaction is 8 Å. First, 3000 steps steepest descent followed by 2000 steps conjugate gradient minimization are performed. After that, additional 7000 steps steepest descent followed by 3000 steps conjugate gradient are performed without restraint. After the minimization, the system is heated up to 300 K within 20 ps. The system is then equilibrated in the NPT ensemble for 100 ps, followed by a further equilibration of 100 ps in NPT. The temperature is controlled at 300 K by Langevin dynamics. The pressure is controlled at 1 bar by isotropic Berendsen barostat. Both the temperature and pressure are regulated every 1 ps. The production MD simulation is carried out in a NPT ensemble with the same parameter settings. The ccptraj module was used for the analysis of trajectories ^51^.

### Cell culture and transfection

HA-tagged human TRIM7 was constructed by inserting into the pCMV-HA vector. Flag-tagged norovirus NTPase, SARS-CoV-2 NSP5 and NSP8 were cloned into the pCMW-Tag2B vector. HEK293T cells were cultured in Dulbecco's modified eagle medium (DMEM, Gibco) supplemented with 10% fetal bovine serum, 2 mM glutamine and 100 U/ml penicillin/streptomycin at 5% CO2 at 37°C. Transfection was performed with polyethyleneimine (PEI). Cells cultured in 6-well plates were prepared in serum-free DMEM media. For each well, 3 μg DNA and 9 μg PEI were pre-incubated in 200 μL serum-free DMEM by vertexing for 15 min. The PEI/DNA mixture was then added into cells and incubated for 4 h at 37°C. Serum-free medium with DNA/PEI mixture was then replaced by the complete medium. Cells were then cultured for 24-48 h to express target proteins.

### Co-immunoprecipitation and immunoblot analysis

HEK293T cells were harvested and washed with PBS buffer before lysed in the NP-40 buffer containing 50 mM HEPES pH 7.4, 150 mM NaCl, 1% (v/v) NP-40 supplemented with the protease inhibitor cocktail. The lysate was cleared by centrifugation at 12,000 rpm at 4°C. For co-immunoprecipitation (co-IP), 500 μg of the supernatant was incubated with 2 μg of antibody followed by incubation with 10 μl 50% slurry of Protein A/G agarose (Thermo Fisher Scientific). After extensive washing, the immunoprecipitated proteins were boiled for SDS-PAGE and immunoblot analysis. Primary antibodies including anti-HA (Bioss, bsm-33003M), anti-FLAG (CST, 14793), anti-Myc (Santa Cruz, sc-40) and anti-actin (CST, 4790) were used.

### Ubiquitination assays

HEK293T cells were co-transfected for 24 h with TRIM7-HA, Ubi-Myc and one of these putative Flag-tagged substrate proteins. Cells were treated with 10 mM MG132 (MCE) for 6 h, washed with cold PBS buffer, harvested and lysed with the NP-40 lysis buffer in the presence of protease inhibitor cocktail. The cleared lysates were used for co-IP using anti-HA (Bioss, bsm-33003M) or anti-FLAG (CST, 14793) antibody. The immunoprecipitated proteins were resolved with SDS-PAGE, transferred to nitrocellulose filter membrane, and incubated with primary antibodies of interest for analysis.

### Lentivirus-mediated stable cell line construction

To generate the TRIM7/TRIM7-mut stable expression cell line, HEK293T cells were transfected with pLV-EGFP-Control/TRIM7/TRIM7-mut, VSV-G, and psPAX2 plasmids to generate lentivirus. The supernatant was harvested and concentrated by 4xPEG-8000 solution 48 h post transfection. HEK293T cells were infected with the concentrated virus, supplied with 8 μg/ml Polybrene. After 48 h of infection, stable EGFP-positive cells were isolated by FACS cell sorting.

### Dual-luciferase reporter assay

HEK293T cells (1×104/well, 96-well white plate) stably expressed TRIM7/TRIM7-mut or EGFP were co-transfected with 20-80 ng of indicated expression plasmids or empty vector, and luciferase plasmid cocktail containing 10 ng of IFN-β-firefly-luc plasmid (IFN-β promoter-driven firefly luciferase plasmid) and 5 ng of pTracer-Renilla-luc plasmid (luciferase control plasmid). Cells were stimulated by co-transfection with 10 ng of pcDNA3-RIG-I (2CARD) plasmids, and empty pcDNA3 vector was used as unstimulated control. Lipofectamine 3000 reagent (Invitrogen, L3000001) was used to transfect these plasmids. 24 h post-transfection, the cells were assayed for dual-luciferase assay activities according to the manufacturer’s instructions (Beyotime, RG088S).

### Analysis of phosphorylation

HEK293T cells (1×105/well, 12-well plate) were co-transfected with TBK1-Myc-expressing plasmid, NSP5/NSP8-encoding plasmids and TRIM7-HA expression plasmid. 24 h post-transfection, western blot was used to analyze the cell lysates for S172-phosphorylated TBK1 (CST, D52C2), total TBK1 (anti-TBK1), and indicated protein.

### Degradation assay

For degradation studies, HEK293T cells (1×105/well, 12-well plate) were co-transfected with indicated expression plasmids encoding potential TRIM7 substrates, and increasing amounts of TRIM7-HA plasmid. To block proteasome, cells were treated with 10 μM MG132 (MedChemExpress, HY-13259) or DMSO (Control) at 18 h post-transfection for 6 h. Then, cells were assessed by western blot.

## Acknowledgements

We thank all the members of Zhang Lab for their kind help. We thank Dr. Jian Lei for supporting plasmids encoding SARS-CoV-2 proteins. We thank the staffs from BL17B/BL18U1/BL19U1/BL19U2/BL01B beamlines of the National Facility for Protein Science in Shanghai (NFPS) at Shanghai Synchrotron Radiation Facility, for assistance during data collection. This work is supported by the National Natural Science Foundation of China Grants (32071218).

## Author contributions

X.L., Y.W., Y.M. and Y.Z. performed the sample preparation, biochemistry and crystallization experiments. Xu.L. collected the diffraction data and solved the structure with the help from X.L. and Y. M.. X.C. performed the Molecular dynamics analysis. J.X., X.L., X.Z., D.J. and P.W. carried out the cellular experiments. C.X., L.L., G.Y. and H.Z. analyzed the data. H.Z., G.Y. and L.L. designed the project and wrote the manuscript.

