## Supplemental info for "Structural insights into the viral proteins binding by TRIM7 reveal a general C-terminal glutamine recognition mechanism"

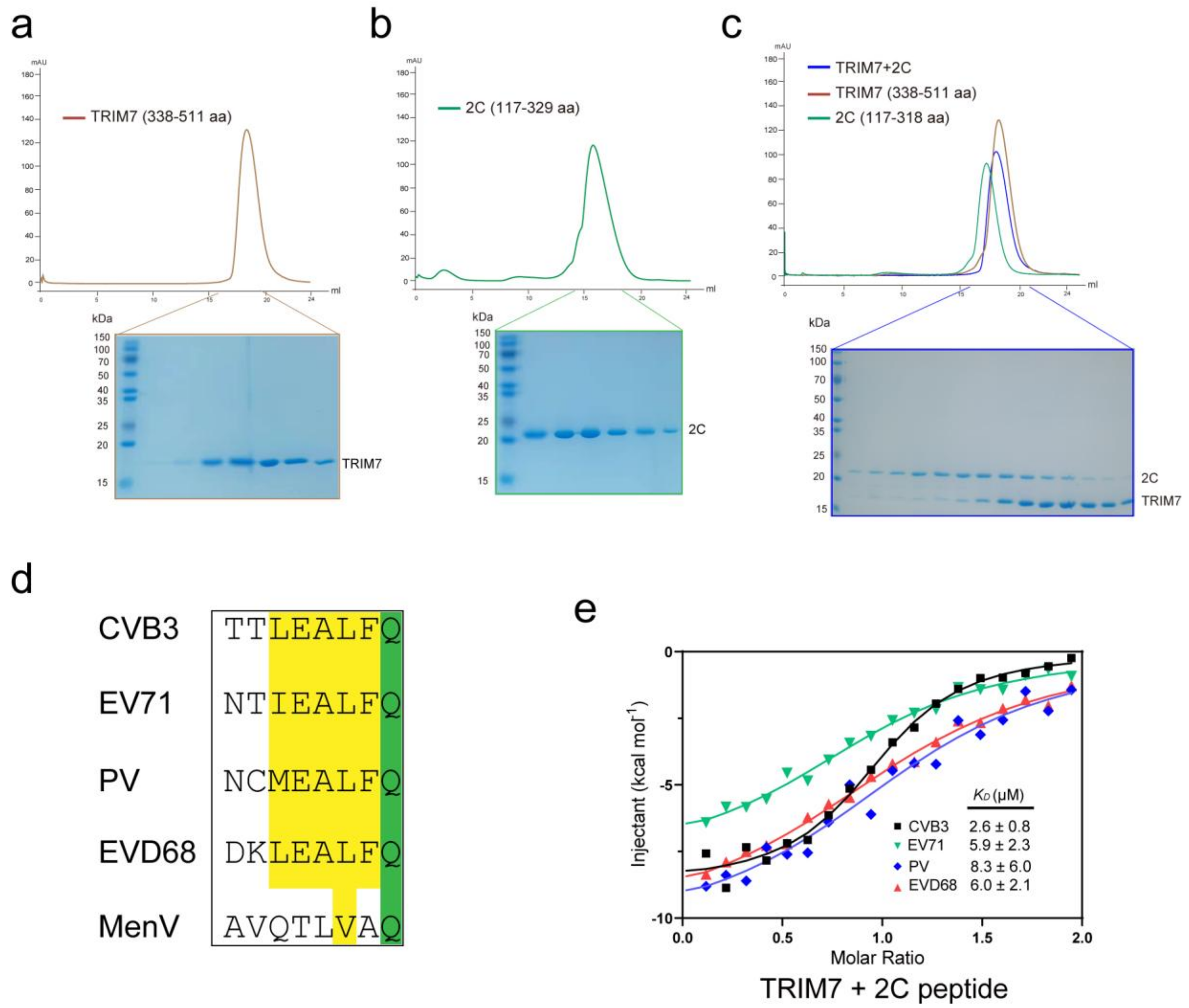


**Figure S1. TRIM7 interacts with enteroviruses proteins.**

**a-b,** SDS-PAGE analysis of the SEC elution fractions corresponding to TRIM7 (a) or 2C (b). **c,** Gel filtration analysis of TRIM7 with C-terminus truncated 2C (117-318 aa). Top panel: elution profile. Bottom panel: SDS-PAGE analysis of elution fractions. **d,** Sequence alignment of 2C proteins from different enteroviruses. Invariant and conserved residues are shaded green and yellow, respectively. **e,** Integrated heat plots for ITC measurements of peptides derived from the C-terminal fragments of 2C proteins from different enteroviruses including CVB3 (black, square), EV71 (green, inverted triangle), PV (blue, rhombus) and EVD68 (red, triangle), to TRIM7.

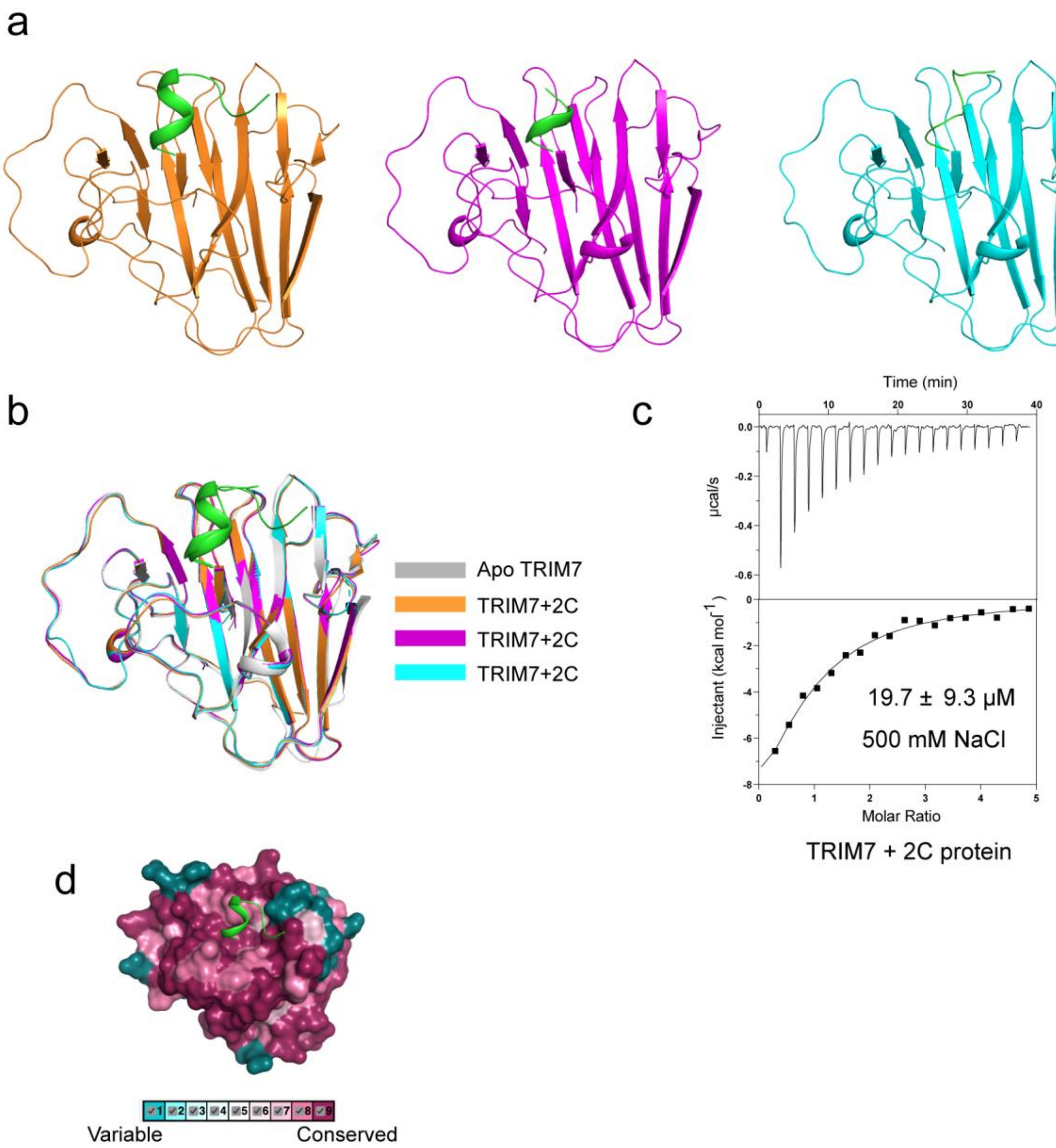


**Figure S2. Structural features of TRIM7-2C complex. a,** Three structures were solved. **b,** Structural comparison of the three TRIM7-2C structures and a previously published substrate-free structure (PDB ID: 6UMA). **c,** ITC measurement of TRIM7-2C association in a high salt buffer containing 500 mM NaCl. **d,** Conservation scores across different species were mapped to the TRIM7 structure.

a

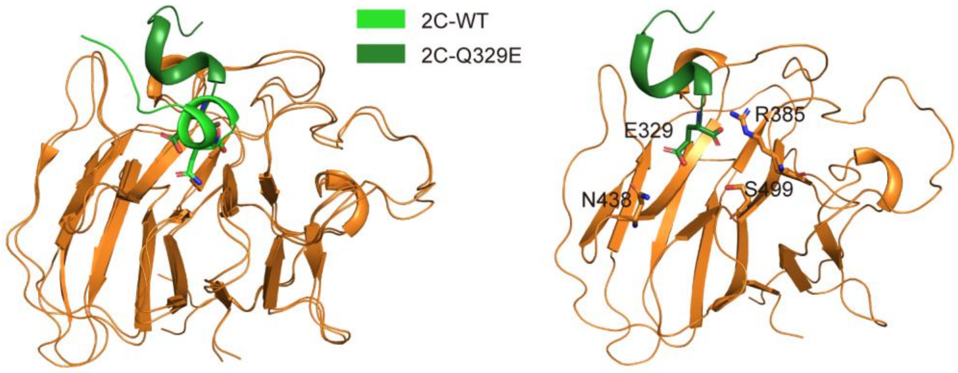


b

Summary of ITC results on TRIM7 interaction with 2C variants

| 2C variants | Kd (μM) |
|---|---|
| TTLEAAFQ | 7.1±1.1 |
| TTLEATFQ | 8.0±1.6 |
| TTLEAVFQ | 8.0±1.3 |
| TTLEAMFQ | 4.1±0.2 |
| TTLEAFFQ | 7.1±1.8 |
| TTLEANFQ | 17.7±1.5 |
| TTLEAEFQ | 9.1±1.0 |
| TTLEARFQ | 11.3±1.4 |

**Figure S3. MD Simulations and ITC measurements of the interaction between TRIM7 and 2C variants.**

**a,** (left) Structural comparison between TRIM2-2C and the calculated TRIM7-2C (Q329E). (right) Effects of simulations on the Q329E mutation. Molecular dynamics simulation suggested that Q329E mutation eliminates the interactions between the residue 329 and TRIM7, especially the interactions with Arg385, Asn438, and Ser499 in TRIM7 (**Table S2**). Consequently, the C-terminal carboxylate group of 2C at residue 329 is more exposed to solvent, leading to a further reduced TRIM7-2C interaction and increased 2C C-terminal solvation. **b,** Summary of ITC results for TRIM7 with 2C variants containing diverse mutations at position -3.

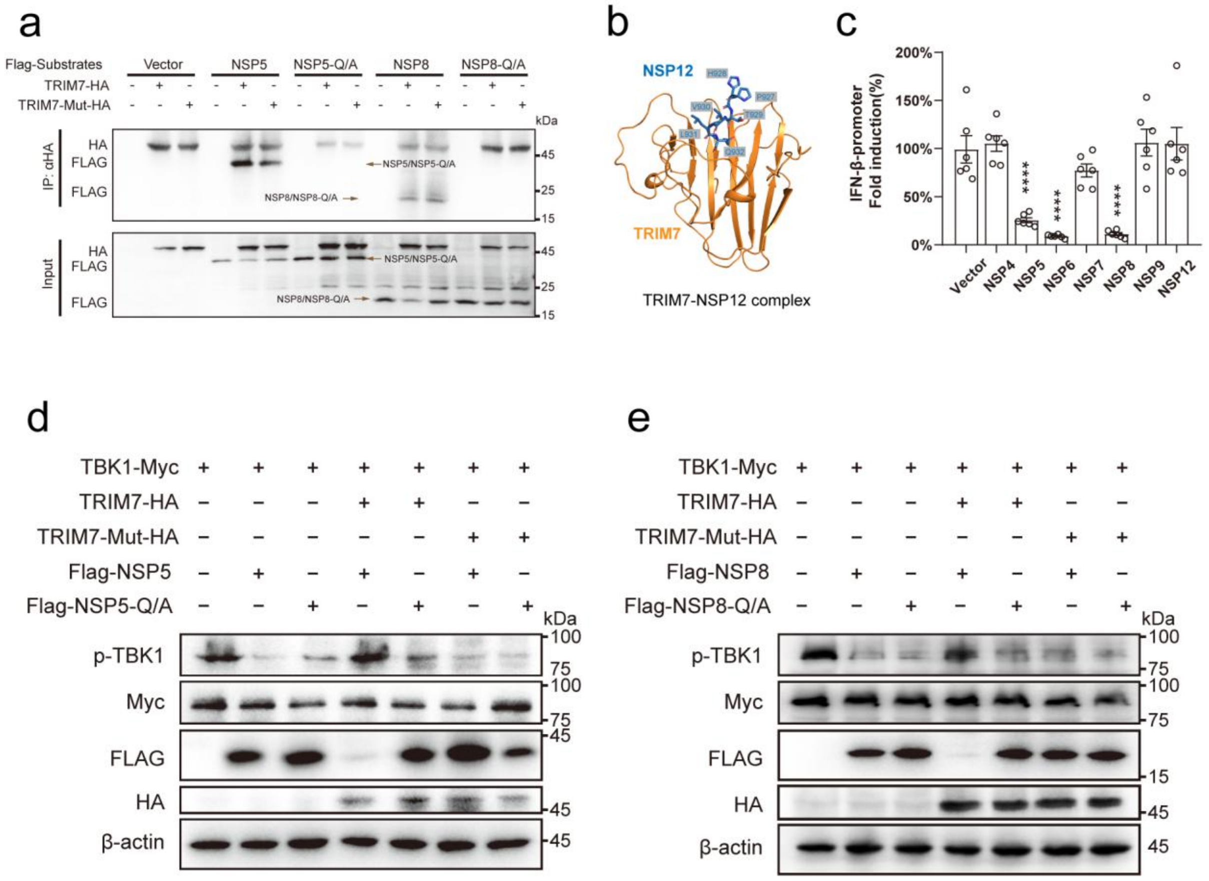


**Figure S4. NSP proteins from SARS-CoV-2 counteract the IFN-β response.**

**a,** Co-immunoprecipitation experiments with NSP5 and NSP8. Flag-tagged NSP5, NSP8 or empty vector was co-transfected with TRIM7-HA in HEK293T cells. Cells were lysed and immunoprecipitated with anti-HA antibody and detected by anti-Flag antibody. TRIM7-Mut-HA, TRIM7 catalytic dead mutant (C29A/C31A); NSP5/NSP8-Q/A, the alanine replacement of the C-terminal glutamine in NSP5/8. **b,** Crystal structure of TRIM7 in complex with NSP12 (residues 928-932). **c,** Luciferase activity of IFN-β promoter reporter in HEK293T cells transfected with indicated NSP proteins and RIG-I CARD expression plasmids. Relative luciferase activity was quantified 24 h post-transfection. Error bars represent mean ± SEM of technical triplicates. Statistical significance was determined by comparing "Vector" group using two-way ANOVA with Dunnett's correction, ****$p < 0.0001$. **d-e,** Analysis of the TBK1 phosphorylation in the presence of NSP5 (d) or NSP8 (e).

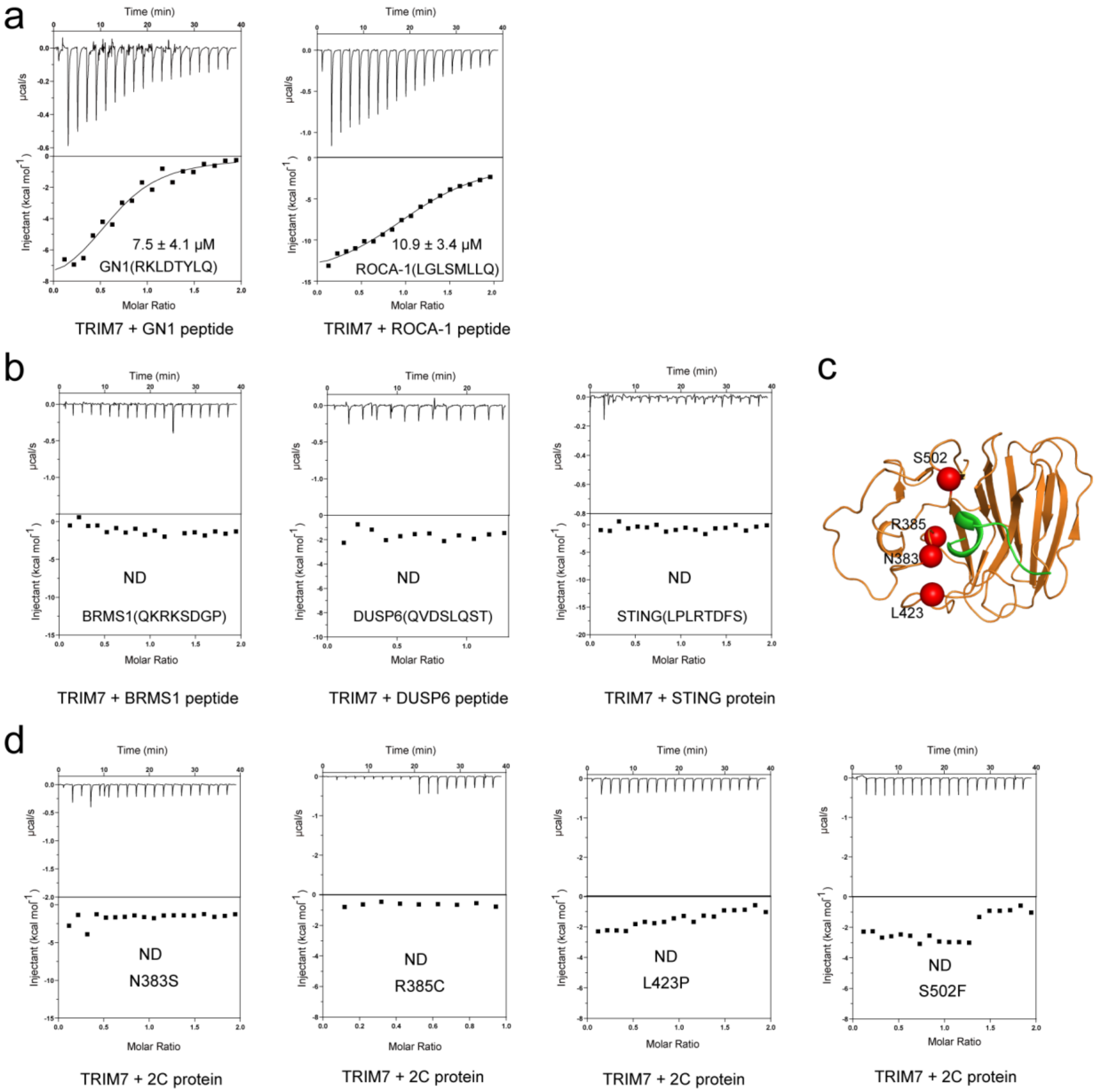


**Figure S5. Binding measurements with multiple proteins functionally linked to TRIM7.**

**a,** TRIM7 binds with GN1 and RACO-1 C-terminus peptide fragments. **b,** Thermodynamic analysis of the interaction between TRIM7 and BRMS1 peptide (left panel), DUSP6 peptide (middle panel) and STING protein (right panel). **c,** Mutations inside the substrate binding pocket of TRIM7 identified from the COSMIC database. **d,** Cancer-associated TRIM7 mutations impaired substrate binding.

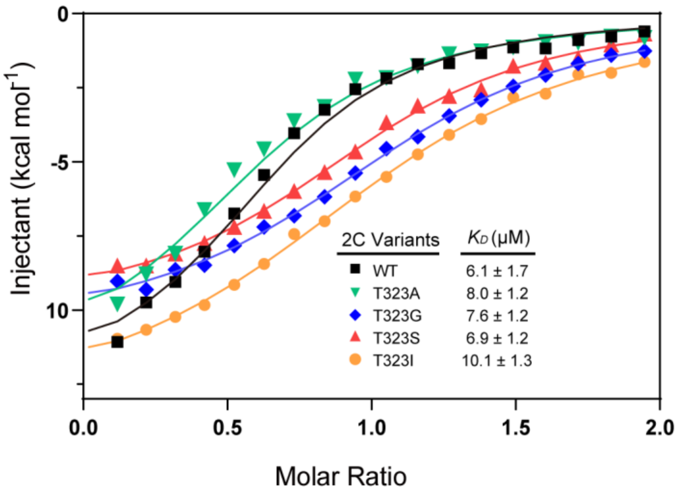


**Figures S6. Quantification of the binding affinity between TRIM7 and CVB3 2C mutations.** Thermodynamic analysis of the interaction between TRIM7 and 2C mutants (T323A, T323G, T323S, T323I).
